# The problem of unmeasured variables in animal social network analysis: can edge-based multilevel models provide a reliable solution?

**DOI:** 10.1101/2024.11.18.623821

**Authors:** Tyler R. Bonnell, Chloe Vilette, Louise Barrett

## Abstract

Recent approaches to analysing social networks suggest that modeling the edges of the network and using multilevel models will produce more informative estimates. These recent methods have been proposed as a way to better handle the dependency structures of social networks, account for biases in data collection, and retain uncertainty when making inferences about social network structures. We find that they have the potential to also effectively handle unmeasured variables that act as statistical confounds in social network analysis. Using simulated data, we highlight that static social network analyses can be used to identify patterns in social networks, but generally cannot be used to identify the underlying mechanisms behind the patterns. To identify mechanisms, we show that taking a dynamic approach and using edge-based models with additive and multiplicative random effects provides a means to identify mechanisms even when statistical confounds are present. Additive and multiplicative random effects also provide information about social structures not captured by the predictors, facilitating exploratory analysis. We suggest that a keep-it-maximal approach for random effects structures is beneficial for edge-based multilevel models of social networks, and that such approaches can be particularly effective when there are unmeasured variables that are not captured by model predictors.

## Introduction

One of the main strengths of social network analysis (SNA) is its ability to capture and describe the interdependence between individuals, and how that interdependence is structured at the level of the group. These interdependencies shape collective behaviors, enabling the identification of emergent properties within a group, such as social cohesion. The structural measures that are extracted from networks (e.g., degree, eigenvector centrality, density, …, etc.), however, can also be highly interdependent themselves as they describe related or overlapping aspects of the network’s structure. This interconnectedness poses a challenge (Webber, Schneider & Vander Wal 2020), especially when attempting to identify causal mechanisms driving changes in social behaviour – whether this entails an attempt to understand how an individual’s position within a social network affects their behaviour, or conversely, how an individual’s behaviour influences their position within the social network.

Although SNA often relies on network patterns extracted directly from networks that have been constructed using observed behavioural interactions (Croft, James & Krause 2008; Krause, Lusseau & James 2009), this direct approach may not fully capture the uncertainty around the way that these relationships (edges) influence the overall network’s structure. It is also often difficult to account for sampling biases in traditional SNA analyses. To address these limitations, there have been calls for an edge-based approach that models the dependencies between edges prior to constructing the network from which patterns are then measured (Hoff 2021; Hart *et al*. 2022; Ross, McElreath & Redhead 2022; Redhead, McElreath & Ross 2023). That is, edge-based models estimate the relationships (edges) between nodes probabilistically, rather than assuming that these edges have very little or no uncertainty. By explicitly modeling edges, this approach can capture the uncertainty in the relationships and account directly for biases in sampling effort (Hart *et al*. 2022).

Here, we aim to test how well edge-based methods perform under realistic conditions, where unmeasured variables may be responsible for shaping individual interactions within a social network (Farine 2024). Unmeasured variables are a common occurrence in animal social networks that can influence social interactions (Holekamp *et al*. 2012; Franks *et al*. 2020), including the presence of shared matrilines, aspects of habitat use within a group’s home range (e.g., similar sleep trees, favourite feeding sites), as well as particular individual characteristics. It is not usually possible to identify and account for all these variables. Furthermore, these unmeasured variables also severely limit our ability to infer the causal mechanisms behind observed patterns in social networks. For instance, if a high level of reciprocity is observed in a network, it is not necessarily an indication that individuals are actively choosing their interactions based on reciprocity. This pattern could instead result from an unmeasured variable that confounds our estimates (VanderWeele & An 2013). Edge-based models, however, open the possibility that we may be able to account for the influence of unmeasured variables through the use of random effect terms in a multilevel modeling framework. Random effects account for unobserved group-level variables that are not directly measured but can be estimated during the model fitting procedure. When included in a model, random effect variables help capture some of the complexity and interdependence arising from these unmeasured influences (Hoff 2021). By accounting for these unmeasured variables, edge-based multilevel models could facilitate the transition from simply measuring network patterns to uncovering the underlying mechanisms that drive these patterns.

Here, we use simulated data to test whether multilevel edge-based models are able to correctly identify the mechanisms that underly the generation of social network patterns. Specifically, we focus on how edge-based models perform when faced with increasing levels of random effect structuring. By testing models with varying random effect structures, we aim to clarify how random effects can mitigate the limitations of missing key variables. These random effect variables can be introduced in various ways, each differing in the extent of interdependence that they are capable of capturing.

To illustrate the impact of unmeasured variables on the results of SNA, we used an example of simulated data that included an unmeasured confounding variable. This unmeasured confounder has an influence on the probability that individuals will interact, and therefore the overall network structure. However, because this variable is unmeasured, we cannot account for it directly in the model. In this context, we compare edge-based approaches to common methods of SNA in animal behaviour to highlight the limitations and strengths of each approach when an unmeasured variable is present. Ultimately, our findings suggest that, to understand the mechanisms driving observed patterns in social behaviour within networks, we need to consider temporal network approaches, which account for changes in network structure over time, and use edge-based models with additive and multiplicative random effects.

## Methods

### Simulating a social network dataset

To simulate social network data, we take a longitudinal approach, which assumes that past interactions will influence future ones. From an animal behavior perspective, this offers a realistic example, as past affiliative and agonistic patterns, especially reciprocity and stability, have been shown to be important mechanisms driving social structure (Antal, Krapivsky & Redner 2005; Puga-Gonzalez *et al*. 2018).

We simulate a group of 20 individuals and allow each individual’s past grooming choices to influence their future grooming choices. Specifically, individuals are more likely to groom their previous grooming partners. In the simulation, the probability of grooming another individual in subsequent networks is based on their interactions in the previous 10 networks. The simulation consists of 100 rounds of interactions, starting with an initial random network of past interactions. Each individual is assigned a specific grooming probability, which is determined using a beta distribution (alpha=5, beta=2), resulting in a population mean probability of grooming of 0.72 at each round of interaction (95%CI: 0.41, 0.97). We also introduce an unmeasured variable that increases the probability of grooming within certain subsets of the group. Specifically, this variable raises the grooming likelihood between individuals 1-4 by 0.02, as well as within the groups 5-15 and 16-20. This adjustment makes individuals within these subgroups slightly more likely to select each other as grooming partners, compared to those outside their subgroups. This unmeasured variable can represent any driver of social interaction, such as shared space use, shared matrilines, or shared sleep sites. The inclusion of this unmeasured factor is meant to capture processes that provide opportunities or constraints on social interactions between subsets of the group and may not be directly observable or measured by a researcher.

Using these simulated data, we assess whether SNA approaches can accurately estimate the correct mechanism driving observed social structures. In this case, the correct underlying mechanism is that past out-grooming interactions alone influence future out-grooming choices, with no influence from alternative social structures when considering whom to groom. Here, we use reciprocity and transitivity as alternative grooming structures that a researcher might test to determine their influence on grooming behaviour. Figure 1 shows the important variables considered in this analysis, highlighting how they are causally interrelated.

**Figure 1:**
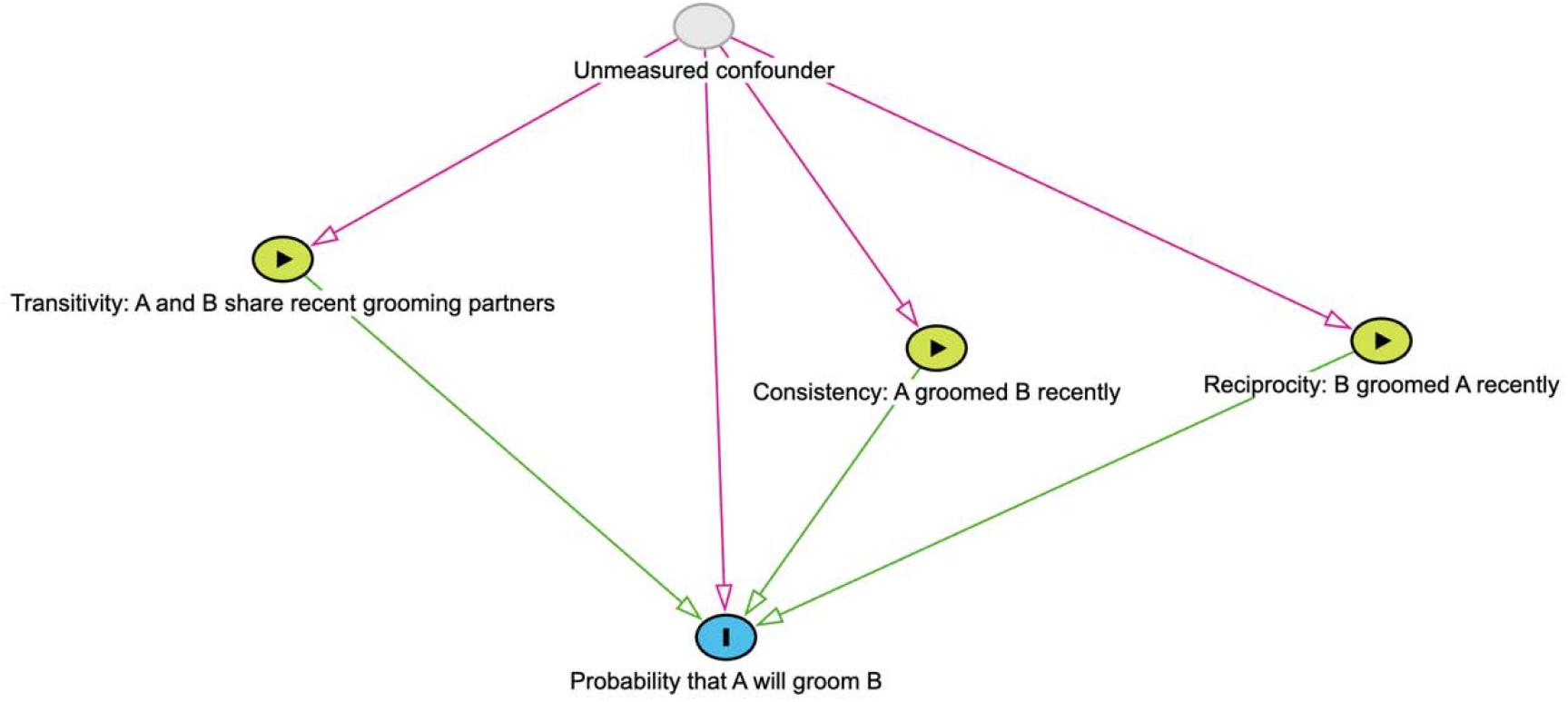
Causal diagram of the potential influences of past interactions and unmeasured variables on the probability of interactions. In the simulated data, only consistency and the unmeasured confounder have any impact on the probability that A will groom B.

In the supplementary section, we perform the same tests on two additional simulated datasets. First, we examine a random grooming dataset, where there was no influence of past interactions on subsequent grooming interactions. In this scenario, grooming choices are entirely random. Second, we use a no-confounder dataset, where past out-grooming choices influence future out-grooming choices, but without an unmeasured confounder variable. In the latter case, the only factor driving grooming choices is the individual’s own previous grooming behavior, without any hidden variable influencing certain sub-groups within the network.

### Social network approaches

We use four different SNA approaches to determine whether these can accurately identify the underlying mechanisms driving observed patterns in social interactions (Table 1). These approaches are categorized into *static* and *dynamic* methods, each with an *observational* and a *modeling* component. While the *dynamic modeling* approach is the only method that directly addresses the question of which social structures influence grooming behaviour, we include the other three methods as they are widely used in SNA. By doing so, we wish to highlight their limitations when it comes to making inferences about the underlying mechanisms driving observed patterns in social networks.

**Table 1:**
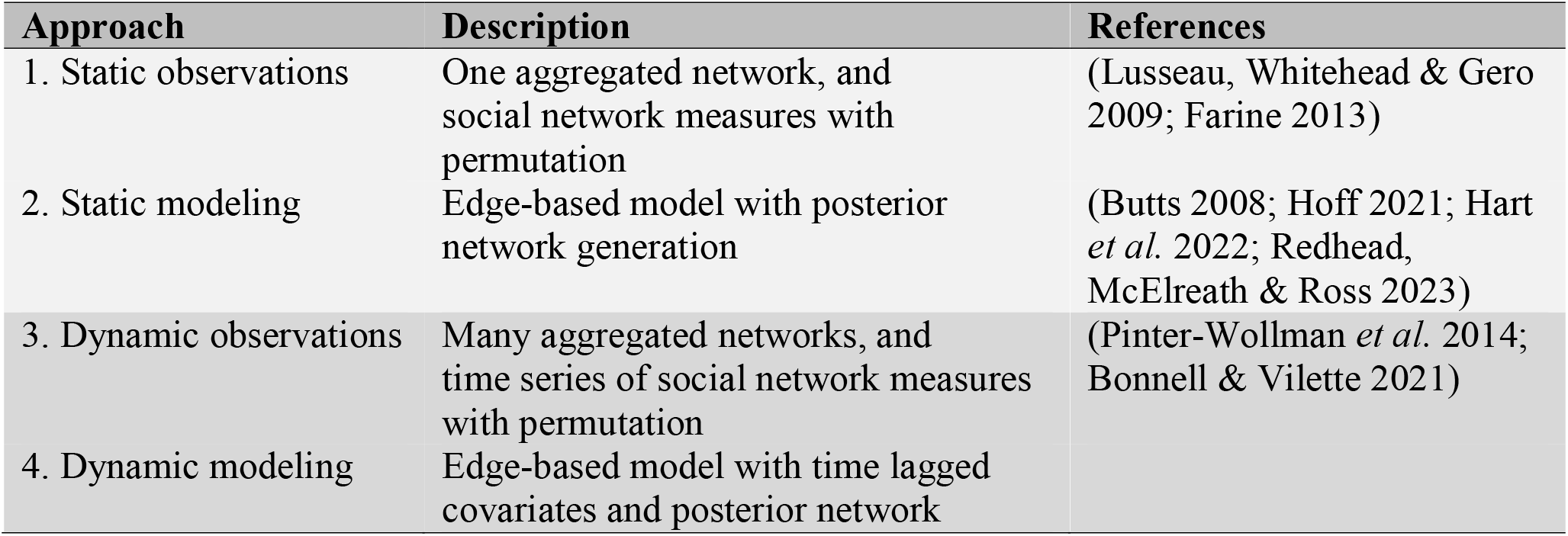
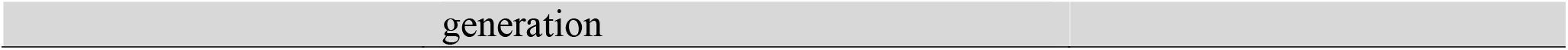
Social network approaches tested.

| Approach | Description | References |
| --- | --- | --- |
| 1. Static observations | One aggregated network, and social network measures with permutation | (Lusseau, Whitehead & Gero 2009; Farine 2013) |
| 2. Static modeling | Edge-based model with posterior network generation | (Butts 2008; Hoff 2021; Hart *et al.* 2022; Redhead, McElreath & Ross 2023) |
| 3. Dynamic observations | Many aggregated networks, and time series of social network measures with permutation | (Pinter-Wollman *et al.* 2014; Bonnell & Vilette 2021) |
| 4. Dynamic modeling | Edge-based model with time lagged covariates and posterior network |  |

The *static* approach relies on aggregating observed social interactions into a single static network and then extracting social network measures. This was used in two ways:

1. A *static observations* approach, which performs pre-network permutations to estimate whether the observed social network measures fall within the range expected if interactions were random (Farine 2017).
2. A *static modeling* approach, where we apply edge-based models to the aggregated network, including increasing levels of random effect structures to account for potential unmeasured variables. Using a Bernoulli model structure, we fit edge-based models to estimates the probability that two nodes will interact. Once the model has been fitted, we use the BISON method to generate a distribution of networks that can be used to make inferences about social network measures, such as reciprocity and transitivity (Hart *et al*. 2022). Our *dynamic* approach (Table 1) differs from the above by accounting for temporal changes in social interactions, allowing us to investigate how past interactions might influence future ones. This approach was again used in two ways:
3. A *dynamic observations* approach that uses a moving window. This method aggregates social interactions over time to produce a time series of measures of social network structure (Bonnell & Vilette 2021). Similar to the *static observations* approach, we use pre-network permutations to assess whether the measured structure (i.e., reciprocity and transitivity) falls within the range expected if grooming interactions were random.
4. A *dynamic modeling* approach, which uses edge-based models with increasing levels of random effect structures. This approach incorporates time-lagged covariates, allowing us to quantify if past social structures can predict future interactions. We fit Bernoulli models with time lagged covariates to estimate the probability that two nodes will interact.

### Statistical models

To analyse our simulated datasets, we use a range of edge-based models, which estimate the probability that two individuals will interact. These models frequently use multilevel modeling frameworks, a common approach in ecological studies (Bolker *et al*. 2009). These edge-based models have been widely studied, especially in the form of social relations models (SRM) (Butts 2008). SRM models allow for the estimation of individual level differences in sending and receiving ties, and also quantifies the reciprocal relation between individuals. Additionally, some edge-based models take a Bayesian approach to directly estimate edge weights, which captures the uncertainty in dyadic relationships (e.g., BISON & STRAND) (Hart *et al*. 2022; Redhead, McElreath & Ross 2023). By directly estimating edge weights, these models facilitate controlling for sampling effort and other covariates that might influence dyadic relationships. Once fitted, these edge-based models can then be used to generate a distribution of networks, thereby allowing the uncertainty in edge weights to propagate forward to network-level analyses, such as estimating uncertainties around social network measures.

While these edge-based models are effective for estimating dyadic probabilities, they may not always capture the dependencies found in many complex network structures. To address this, methods like stochastic block modeling (Lee & Wilkinson 2019; Redhead, McElreath & Ross 2023) have been developed. Stochastic block models group individuals into shared “blocks” or clusters, where individuals within the same block have a higher probability of interacting with each other. Similarly, latent space models estimate the positions of individuals within a latent space, where those that are closer together have a higher probability of interacting (Hoff, Raftery & Handcock 2002). More recently, latent factor models (Additive and Multiplicative Effects Network: AMEN) have been introduced as a method that can capture network structures similar to both the stochastic block and latent space models (Hoff 2021).

If we look at these models together, we can see a progression in the ability of the models to capture more social structure through the use of random effect structures. If we assume a Bernoulli distribution with a logit link function for the probability that individual i and j will interact (p _*i,j*_), we can see that each model can be written as an extension of the others with additional random effect structures:

BISON

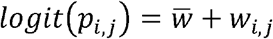

BISON & SRM

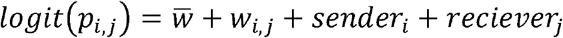

AMEN

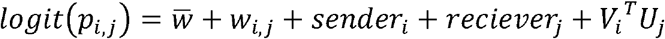

The BISON method estimates the edge weight between sender *i* and receiver *j* (*w*_*i,j*_), as well as the covariance between reciprocal edges (e.g., if A interacts with B, this may influence B interacting with A). The BISON SRM model adds in node-level sender and receiver random effects (*sender*_*i*_ + *reciever*_*j*_), and the covariance between sender/receiver nodes. Finally, AMEN adds in the potential for multiple edge structures (e.g., transitivity, clustering) by adding multiplicative random effects for each node (*V*_*i*_^*T*^ *U*_*j*_).

To further expand this range of models, we add a standard multilevel model (MLM). This is the simplest model in our progression of models. It includes random intercepts for directed dyad IDs, as well as sender and receiver effects, which account for individual differences in tendencies to interact or be interacted with, but has no covariance structures between nodes in a dyad, or between senders and receivers. Finally, we introduce one additional model: the regularized additive and multiplicative effects network model (RAMEN). This model prioritizes random effect variables over predictor variables by applying a weakly informative prior (centered on zero) on the sum of the effects of the predictors. The reasoning behind this prior is to encourage the model to focus on capturing structural dependencies of edges in the random effects.

Taken together, the progression of five models, from MLM to RAMEN, allows us to incorporate increasing levels of social structure into the random effects, thereby capturing increasingly complex network dependencies.

The MLM models were run using brms (Bürkner 2017), while the remaining models were run directly from stan code (Carpenter *et al*. 2017). All model code is available on GitHub: https://github.com/tbonne/AMEN_BISON.

### Comparing outcomes

With our simulated data we know the correct outcome to expect, namely that only past out-grooming should influence grooming preferences, with an unmeasured confound making it more likely for certain sub-groups to groom each other.

For the *static observation* approach, we use the walktrap algorithm and hierarchical clustering in igraph package of R (Csardi & Nepusz 2006) to determine whether the known sub-groups are detected. We also perform pre-network permutations on reciprocity and transitivity measures on 100 simulated datasets.

For the *static modeling* approach, we fit the edge-based model and generated networks from the fitted models using posterior network predictions. We then used those posteriors networks to estimate reciprocity and transitivity in the network. The RAMEN model was excluded in these tests, as no predictor variables are used in our static modeling approach. Furthermore, to assess how well the networks generated from model predictions aligned with the true social structure underlying the simulated data, we compared clustering in the posterior networks with the true networks. This involved comparing networks generated from model predictions (N=100) with the true network structure set during simulation. We estimated the similarity in community clustering between the simulated and true networks using the adjusted rand index from the mclust package in R (Scrucca *et al*. 2016). Clustering was determined using the walktrap algorithm.

For the *dynamic observation* approach, we plotted the simulated data over time by aggregating the social networks into time windows, and extracting reciprocity and transitivity measures. We perform pre-network permutations to these data to determine what measures are outside the range of chance interactions. We perform these steps on 100 simulated datasets using the netts package in R (Bonnell & Vilette 2021).

Finally, for the *dynamic modeling* approach, we fit the edge-based models and use the time-lagged effects to determine which past social structures are likely to be influencing future grooming behaviours. We record whether the model successfully identifies past out-grooming as an influential variable, while identifying that both reciprocity and transitivity are not influential. We used the 95% credible interval to determine if zero was a likely value. We ran these models on 100 simulated datasets and plot the percentage of times that the models correctly identified the correct influential variables.

## Results

### Static observation approach – working directly with observations

When analysing the aggregated simulated data, we found that a walktrap algorithm and hierarchical clustering correctly identified the sub-groups within the network (i.e., individual 1-4, 5-15, 16-20) (Fig. 2). When we applied pre-network permutations to 100 simulated datasets, we found that reciprocity and transitivity of the observed network fell outside the range expected by chance in 0.56% and 0.99% of simulations, respectively.

**Figure 2:**
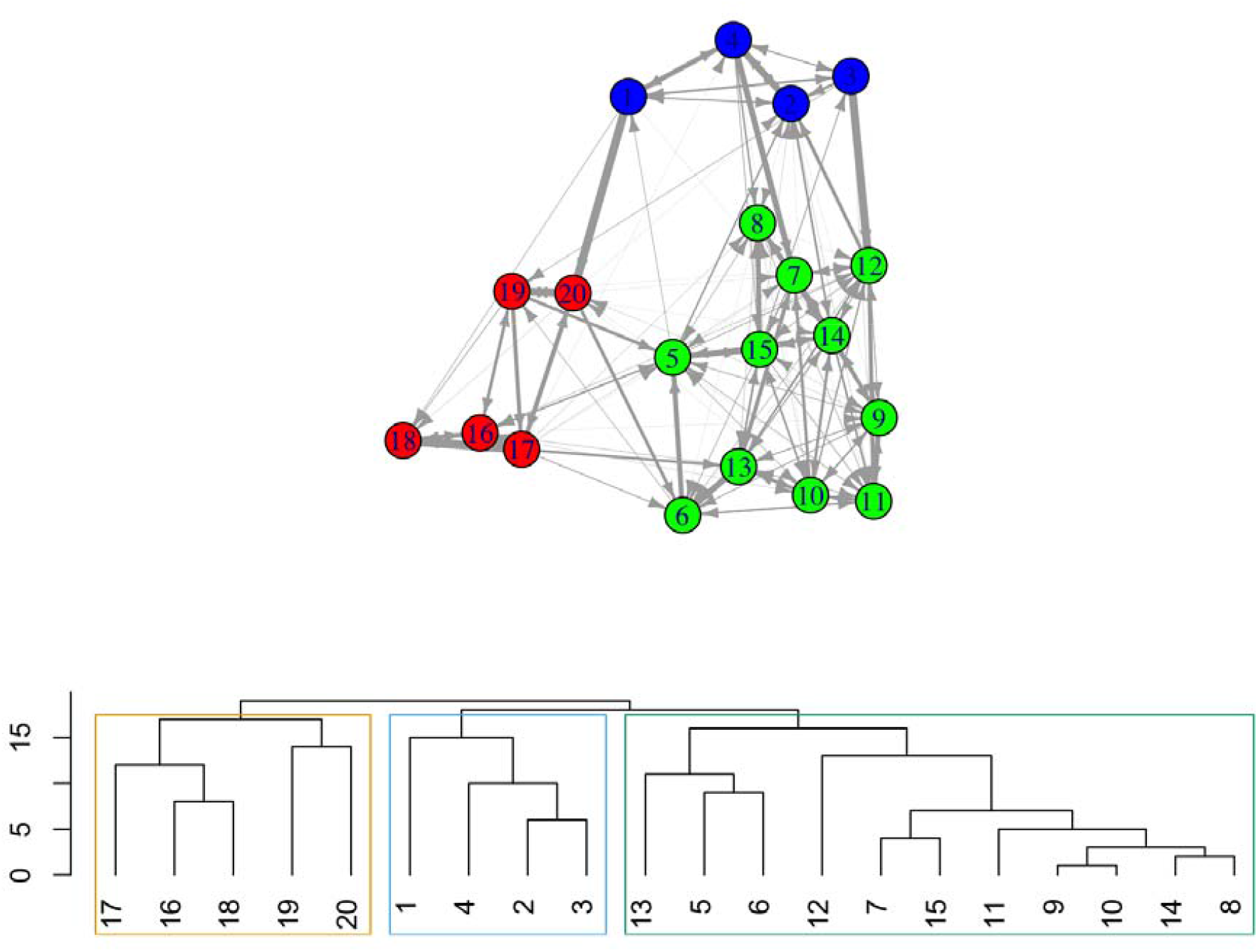
Simulated data aggregated into one network. The walktrap algorithm is used to find communities (colored in the network), while a hierarchical clustering algorithm is used to visualize those clusters. The unmeasured variable increases grooming interactions within subgroups: (1-4, 5-15, 16-20). The only other grooming influence was an individual’s past out-grooming patterns, which makes continuing past grooming choices more likely in the simulated data.

### Static modeling approach – modeling the observations

When we fit edge-based models to the simulated data (i.e., MLM, BISON, BISON SRM, and AMEN), we found that all edge-based models, except MLM, gave similar estimates for patterns of reciprocity and transitivity (Table 1).

**Table 1:**
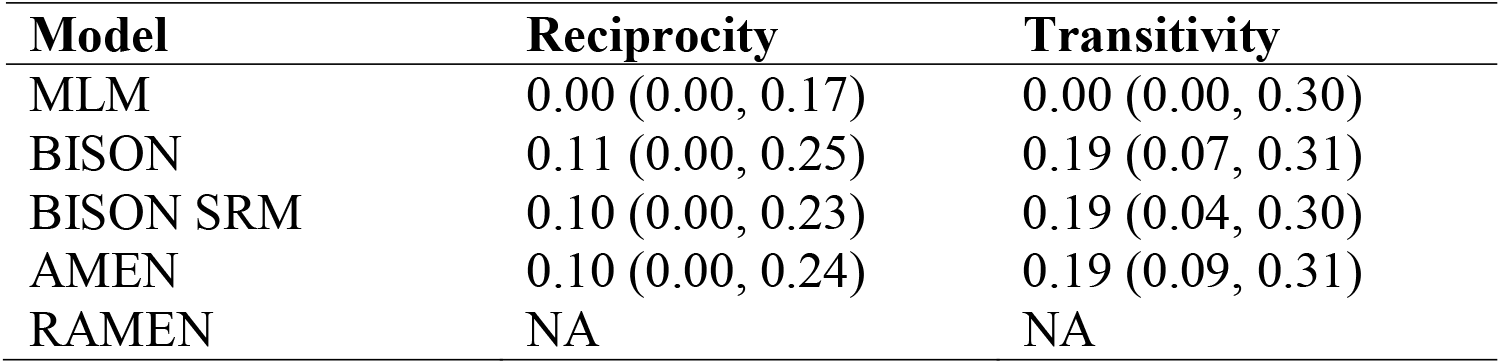
Estimated network characteristics based on posterior networks generated from edge-based models. Median and 95% credible intervals are calculated from 100 model generated networks predicting dyadic interactions during one round of interactions.

| Model | Reciprocity | Transitivity |
| --- | --- | --- |
| MLM | 0.00 (0.00, 0.17) | 0.00 (0.00, 0.30) |
| BISON | 0.11 (0.00, 0.25) | 0.19 (0.07, 0.31) |
| BISON SRM | 0.10 (0.00, 0.23) | 0.19 (0.04, 0.30) |
| AMEN | 0.10 (0.00, 0.24) | 0.19 (0.09, 0.31) |
| RAMEN | NA | NA |

We also found that all edge-based models reproduced the true network structure (Fig. 3).

**Figure 3:**
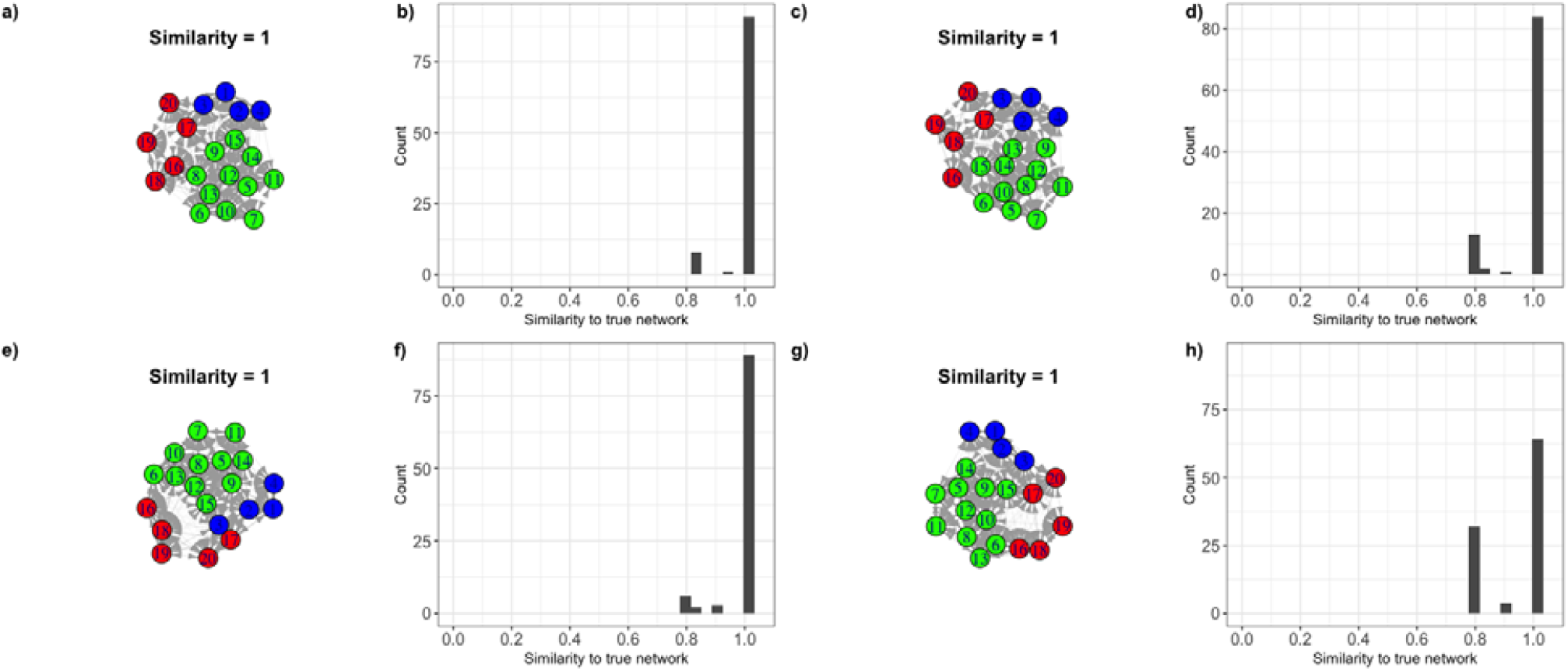
Similarity measures between posterior networks generated by MLM (a, b), BISON (c, d), BISON SRM (e, f), and AMEN (g, h) models. The similarity score is a distance measure (rand index) where the larger the distance, the farther the community detection algorithm was from the true network. The networks show one output of the posterior network checks, and the histogram shows the distribution of similarity scores on 100 model generated networks.

### Dynamic observation approach – Working directly with the observations

Using a dynamic observation approach, we observed that both reciprocity and transitivity were generally lower when compared to random grooming patterns (Fig. 4). When the permutation approach was applied on 100 simulated datasets, we found that 40% (95% CI: 14, 62) of the reciprocity and 89% (95% CI: 68, 100) of the transitivity measures fell outside the range of random grooming patterns.

**Figure 4:**
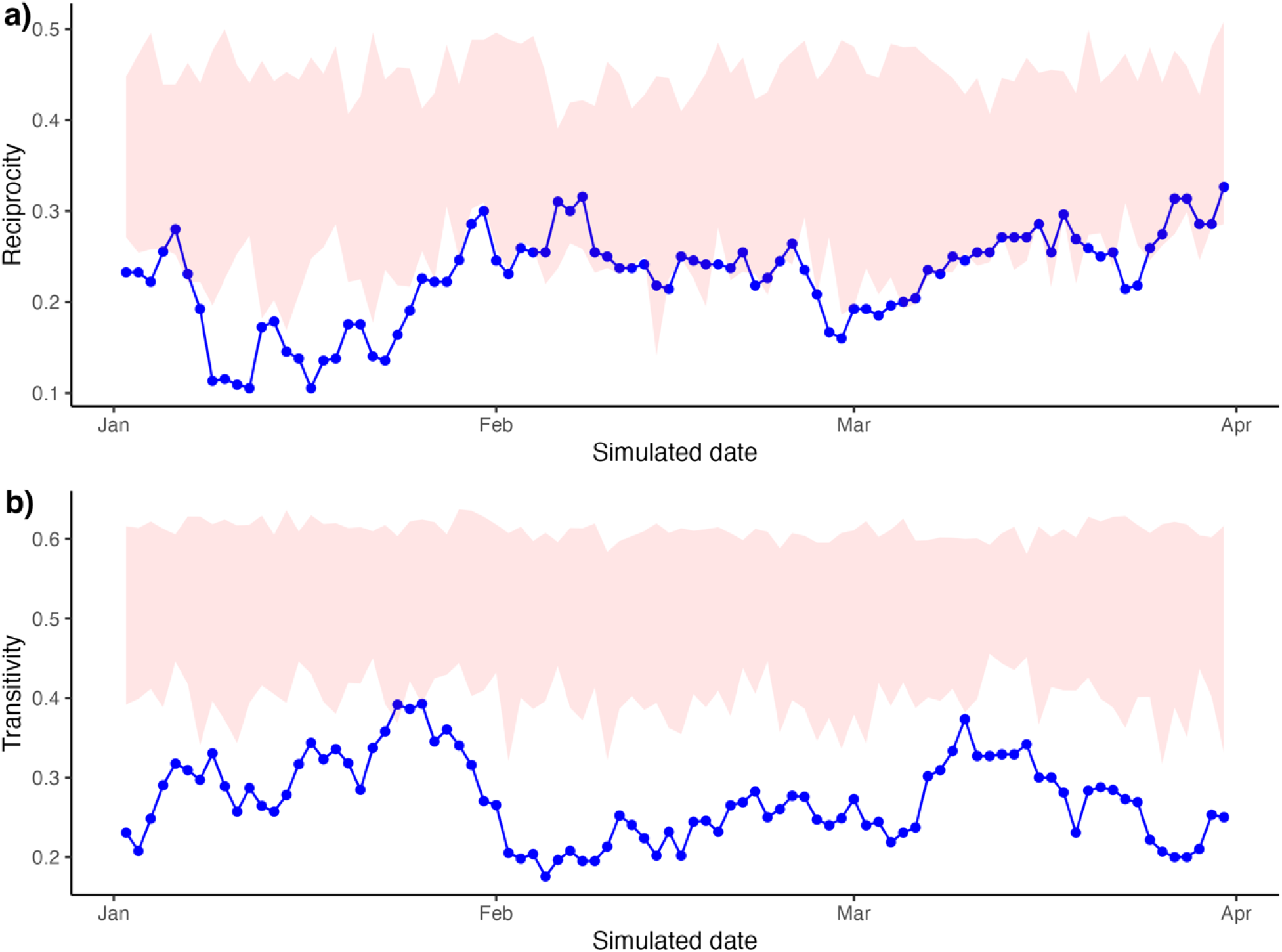
Measures of (a) reciprocity and (b) transitivity through time for one simulated dataset. The pink shaded region represents the region of values likely after 1000 event permutations (i.e., the range expected by chance).

### Dynamic modeling approach – Modeling the observations through time

When using a dynamic modeling approach (i.e., MLM, BISON, BISON SRM, AMEN, RAMEN), all models correctly identified previous out-grooming behaviour as an influential parameter on future grooming behaviour (Fig. 5a). Overall, we found that the models often incorrectly attributed the influence of reciprocity and transitivity on grooming behaviours, when an individuals’ grooming behaviour was, in fact, influenced by a shared, unmeasured variable. Specifically, the MLM model incorrectly identified reciprocity as influential in current grooming behaviours 27% of the time. The error rate dropped for other models, with BISON at 8%, BISON SRM at 9%, AMEN at 14%, and RAMEN at 0% (Fig. 5b). For transitivity, the performance was worse: the MLM model incorrectly estimated transitivity 59% of the time, BISON was incorrect 60% of the time, and BISON SRM was incorrect 53% of the time. The AMEN model performed better, with only 22% of estimates incorrect, while RAMEN had no incorrect estimates (0%) (Fig. 5c).

**Figure 5:**
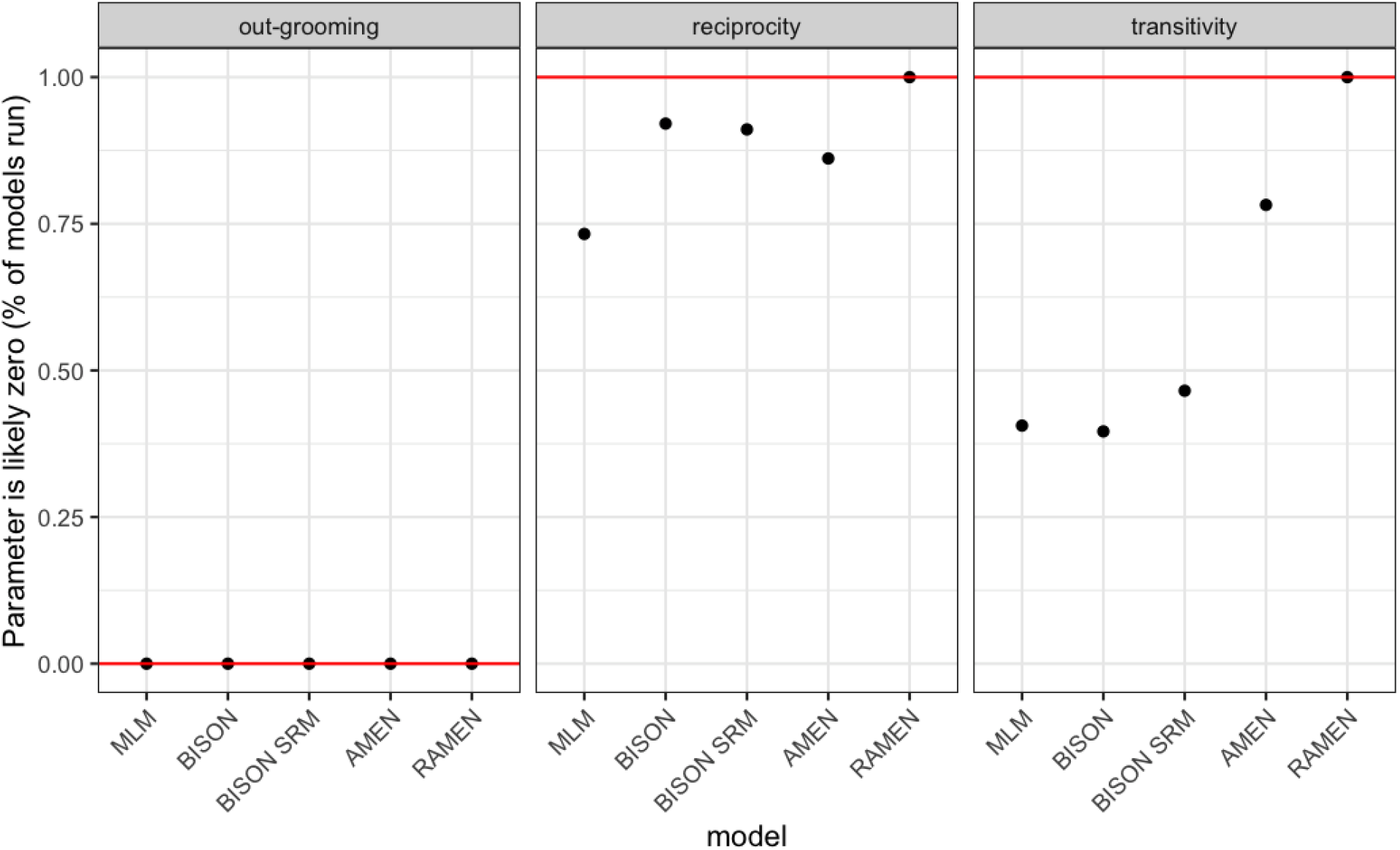
The figure shows the percentage of model runs where the influence of past out-grooming, reciprocity, and transitivity on current grooming patterns was likely zero (i.e., using the 95%CI). The correct answer is represented by the red line. The x-axis shows each model tested, and the y-axis shows the percentage of times that the model found zero to be a credible estimate. New simulated data were generated for each model run.

Results for simulated datasets without a confounder and for datasets where grooming did not depend on past interactions are shown in figures S1 and S2, respectively.

## Discussion

Our results suggest that unmeasured confounding variables present an important challenge for all SNA approaches tested. Nevertheless, the performance of dynamic edge-based models with additive and multiplicative random effect structures suggest it is a promising approach when unmeasured confounds are present.

For the static approaches, we found SNA methods failed to reveal the underlying mechanisms responsible for the observed social structures. By aggregating and extracting measures, either directly or through edge-based models, we identified both transitivity in the network structure and limited reciprocity. This was also the case for the dynamic approach using observations. Therefore, although these methods were able to detect some structural features, they cannot uncover why these patterns exist. Given that we know the network structures were generated by individuals being more likely to groom those they groomed previously, and that tendency was influenced by an unmeasured variable, it would be incorrect and misleading to infer any mechanistic underpinnings from the patterns of reciprocity or transitivity. That is, observed reciprocity in a network does not imply that individuals are tracking reciprocity when interacting with others. Similarly, if transitivity is observed, it does not suggest that individuals are using information about friends of friends when choosing with whom to interact. Another way to say this is that, in our simulated dataset, observed patterns of reciprocity and transitivity resulted from the unmeasured confounder that induced sub-grouping within the population, not from individual choices or strategies. Although this is clear in our simulation, where we know the true underling structuring principles, this is not the case in real-world situations, where confounding factors are often present but difficult to observe or measure, and static approaches may similarly fall short in identifying causal mechanisms behind social structures.

With a dynamic modeling approach using lagged covariates, it became possible to distinguish between mechanism and pattern by using past interactions as predictors of future behaviour. This approach enables the quantification of how previous social interactions influence future ones, allowing for a deeper understanding of social dynamics. We found that the flexibility of the random effects was essential in capturing the interdependencies between network edges and estimating influential variables effectively. When the structure of the random effects was limited, the likelihood of misidentifying variables as predictive of future behaviour increased (Fig. 5). As we enhanced the flexibility of the random effect structure, the models became better at avoiding the misidentification of structured patterns resulting from an unmeasured confounder. The type of statistical confound added in the simulated data is a common one and is known as a fork confound (Pearl 2009; Pearl & Mackenzie 2018). This occurs when a variable is a common cause of two other variables, potentially inducing a spurious relationship between them. In our simulation, the unmeasured confound influenced both the probability of interactions between individuals and the reciprocity and transitivity of the network as a whole. Thus, in scenarios involving such confounds, it is critical to include these confounding variables in model. When these confounding variables are unmeasured, the random effect structures can take on this role, as suggested by our results. Specifically, incorporating additive and multiplicative random effect structures to a model (AMEN) can help account for the interdependencies between edges and, to some extent, address the influence of unmeasured variables. However, even with the AMEN approach, which incorporated the most flexible random variable structure, there were instances where transitivity was misidentified as an important influence on current grooming in about 22% of the simulated cases (Fig. 5). Additionally, AMEN performed slightly worse with reciprocity when compared to BISON or BISON SRM (Fig. 5). When we applied regularization to the model (RAMEN), using a prior that restricted the range of likely values for the total impact of predictors, the incorrect identification dropped considerably (0%). This result suggests that regularized versions of the AMEN could benefit from further investigation, as there may be various ways to prioritize random effect variables over predictors.

An additional benefit of including both additive and multiplicative random effects was the opportunity for exploratory analysis. While the predictors provide estimates of how past interactions predict current behaviour, the random effect variables can estimate the unmeasured structures that remain after accounting for the predictors. This is particularly valuable for exploratory analysis and generating new research questions. For example, one can visualise the position of nodes using the estimated multiplicative random effects, revealing which nodes are closer together and hence more likely to interact (see Fig. S3 for an example).

When estimating network measures, it is beneficial to propagate the uncertainty in the relationships (i.e., edges) by generating networks from the edge-based model predictions and using these networks to extract relevant measures. Our results indicated that all methods tested were able to capture community structure of the simulated data. While no formal tests were conducted, the larger AMEN model exhibited the highest number of cases of misclassification among nodes (Fig. 3h). This suggests that the larger model might require more data than the 100 rounds of interactions to reliably predict discrete communities, revealing a potential trade-off between model flexibility and pattern detection when constructing posterior networks. This trade-off warrants further investigation, as generating accurate posterior networks is crucial for downstream tasks after model fitting in the BISON approach, such as estimating the mean and credible intervals of network measures.

## Conclusion

We demonstrate that static approaches to SNA can only capture structural patterns, making it difficult to make inferences about the underlying mechanisms (Ostner & Schülke 2018). For instance, our simulated data showed that individuals not engaged in reciprocity or influenced by transitivity can still generate these patterns at the network level. In contrast, employing dynamic edge-based modeling approaches to SNA allows for a reliable identification of the mechanisms driving individual social behaviour, particularly when additive and multiplicative random effect structure are included in the models. For example, models with simpler random effect structures misidentified reciprocity and transitivity as important drivers of grooming behaviour. Our results are especially relevant in cases where predictor variables in the model fail to capture all the drivers that provide opportunities or constraints on individuals’ interactions, a highly likely scenario in most animal social networks. Furthermore, our results align with general recommendations for maintaining high flexibility in multilevel models (Barr *et al*. 2013), suggesting that the “keep-it-maximal” approach when choosing random effect structures in multilevel models is also a useful approach in edge-based social network models.

## References

Antal, T., Krapivsky, P.L. & Redner, S. (2005) Dynamics of social balance on networks. Physical Review E—Statistical, Nonlinear, and Soft Matter Physics, 72, 036121.

Barr, D.J., Levy, R., Scheepers, C. & Tily, H.J. (2013) Random effects structure for confirmatory hypothesis testing: Keep it maximal. Journal of memory and language, 68, 255–278.

Bolker, B.M., Brooks, M.E., Clark, C.J., Geange, S.W., Poulsen, J.R., Stevens, M.H.H. & White, J.-S.S. (2009) Generalized linear mixed models: a practical guide for ecology and evolution. Trends in Ecology & Evolution, 24, 127–135.

Bonnell, T.R. & Vilette, C. (2021) Constructing and analysing time-aggregated networks: The role of bootstrapping, permutation and simulation. Methods in Ecology and Evolution, 12, 114–126.

Bürkner, P.-C. (2017) brms: An R package for Bayesian multilevel models using Stan. Journal of statistical software, 80, 1–28.

Butts, C.T. (2008) A relational event framework for social action. Sociological Methodology, 38, 155–200.

Carpenter, B., Gelman, A., Hoffman, M.D., Lee, D., Goodrich, B., Betancourt, M., Brubaker, M., Guo, J., Li, P. & Riddell, A. (2017) Stan: A probabilistic programming language. Journal of statistical software, 76, 1–32.

Croft, D.P., James, R. & Krause, J. (2008) Exploring animal social networks. Princeton University Press.

Csardi, G. & Nepusz, T. (2006) The igraph software. Complex syst, 1695, 1–9.

Farine, D.R. (2013) Animal social network inference and permutations for ecologists in R using asnipe. Methods in Ecology and Evolution, 4, 1187–1194.

Farine, D.R. (2017) A guide to null models for animal social network analysis. Methods in Ecology and Evolution, 8, 1309–1320.

Farine, D.R. (2024) Modelling animal social networks: New solutions and future directions. Journal of Animal Ecology, 93, 250–253.

Franks, V.R., Ewen, J.G., McCready, M., Rowcliffe, J.M., Smith, D. & Thorogood, R. (2020) Analysing age structure, residency and relatedness uncovers social network structure in aggregations of young birds. Animal Behaviour, 166, 73–84.

Hart, J.D.A., Weiss, M.N., Franks, D.W. & Brent, L.J.N. (2022) BISoN: A Bayesian Framework for Inference of Social Networks. bioRxiv, 2021.2012.2020.473541.

Hoff, P. (2021) Additive and Multiplicative Effects Network Models. Statistical Science, 36, 34-50, 17.

Hoff, P.D., Raftery, A.E. & Handcock, M.S. (2002) Latent space approaches to social network analysis. Journal of the american Statistical association, 97, 1090–1098.

Holekamp, K.E., Smith, J.E., Strelioff, C.C., Van Horn, R.C. & Watts, H.E. (2012) Society, demography and genetic structure in the spotted hyena. Molecular Ecology, 21, 613–632.

Krause, J., Lusseau, D. & James, R. (2009) Animal social networks: an introduction. Behavioral Ecology and Sociobiology, 63, 967–973.

Lee, C. & Wilkinson, D.J. (2019) A review of stochastic block models and extensions for graph clustering. Applied Network Science, 4, 1–50.

Lusseau, D., Whitehead, H. & Gero, S. (2009) Incorporating uncertainty into the study of animal social networks. arXiv preprint arXiv:0903.1519.

Ostner, J. & Schülke, O. (2018) Chapter Four - Linking Sociality to Fitness in Primates: A Call for Mechanisms. Advances in the Study of Behavior (eds M. Naguib, L. Barrett, S.D. Healy, J. Podos, L.W. Simmons & M. Zuk), pp. 127–175. Academic Press.

Pearl, J. (2009) Causality. Cambridge university press.

Pearl, J. & Mackenzie, D. (2018) The book of why: the new science of cause and effect.

Basic books.

Pinter-Wollman, N., Hobson, E.A., Smith, J.E., Edelman, A.J., Shizuka, D., De Silva, S., Waters, J.S., Prager, S.D., Sasaki, T. & Wittemyer, G. (2014) The dynamics of animal social networks: analytical, conceptual, and theoretical advances. Behavioral Ecology, 25, 242–255.

Puga-Gonzalez, I., Ostner, J., Schülke, O., Sosa, S., Thierry, B. & Sueur, C. (2018) Mechanisms of reciprocity and diversity in social networks: a modeling and comparative approach. Behavioral Ecology, 29, 745–760.

Redhead, D., McElreath, R. & Ross, C.T. (2023) Reliable network inference from unreliable data: A tutorial on latent network modeling using STRAND. Psychological Methods.

Ross, C.T., McElreath, R. & Redhead, D. (2022) Modelling human and non-human animal network data in R using STRAND. bioRxiv, 2022.2005.2013.491798.

Scrucca, L., Fop, M., Murphy, T.B. & Raftery, A.E. (2016) mclust 5: clustering, classification and density estimation using Gaussian finite mixture models. The R journal, 8, 289.

VanderWeele, T.J. & An, W. (2013) Social networks and causal inference. Handbook of causal analysis for social research, 353–374.

Webber, Q.M.R., Schneider, D.C. & Vander Wal, E. (2020) Is less more? A commentary on the practice of ‘metric hacking’ in animal social network analysis. Animal Behaviour, 168, 109–120.

