## Supplementary material for "The problem of unmeasured variables in animal social network analysis: can edge-based multilevel models provide a reliable solution?"

**Supplementary section**

*Model code*

Full model code written in stan is available on github: https://github.com/tbonne/AMEN_BISON

*Models fit to random grooming data*

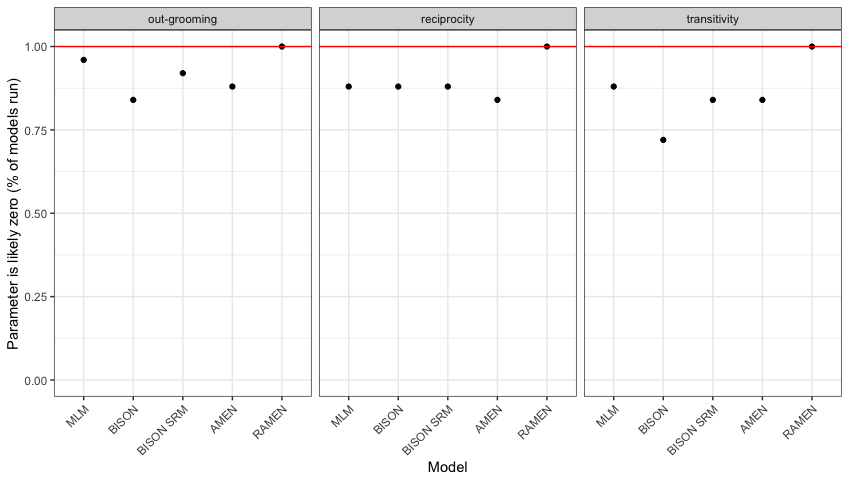

Figure S1: Percentage of models run where the influence of past grooming-out, grooming-in, and transitivity on current grooming patterns was likely zero (i.e., using the 95%CI). The correct answer is represented by the red line, as in these simulated data grooming was not related to past interactions between individuals. The x-axis shows the model used, and the y-axis shows the percentage of times that the model found zero to be a credible estimate (N=30 models).

*Models fit to grooming data where past out-grooming impacts future grooming and no confounding variables.*

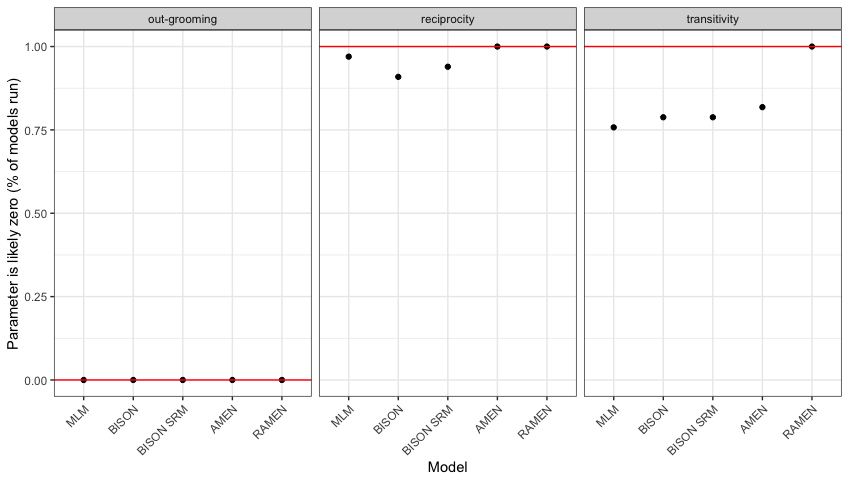

Figure S2: Percentage of models run where the influence of past grooming-out, grooming-in, and transitivity on current grooming patterns was likely zero (i.e., using the 95%CI). The correct answer is represented by the red line, as in these simulated data grooming was influenced by past out-grooming with no other confounding variables present. The x-axis shows the model used, and the y-axis shows the percentage of times that the model found zero to be a credible estimate (N=30 models).

*Unexplained social structures*

Using the AMEN modelling approach, we can see the remaining variation left after accounting for the main effects (Fig. S3). Visualizing this remaining variation can be very useful for exploratory analysis, i.e., we can see the sub-groups clustering: 1-4, 5-15, and 16-20 (Fig. S3), these patterns are not picked up by the main effects in the model but can be seen in the random effects.

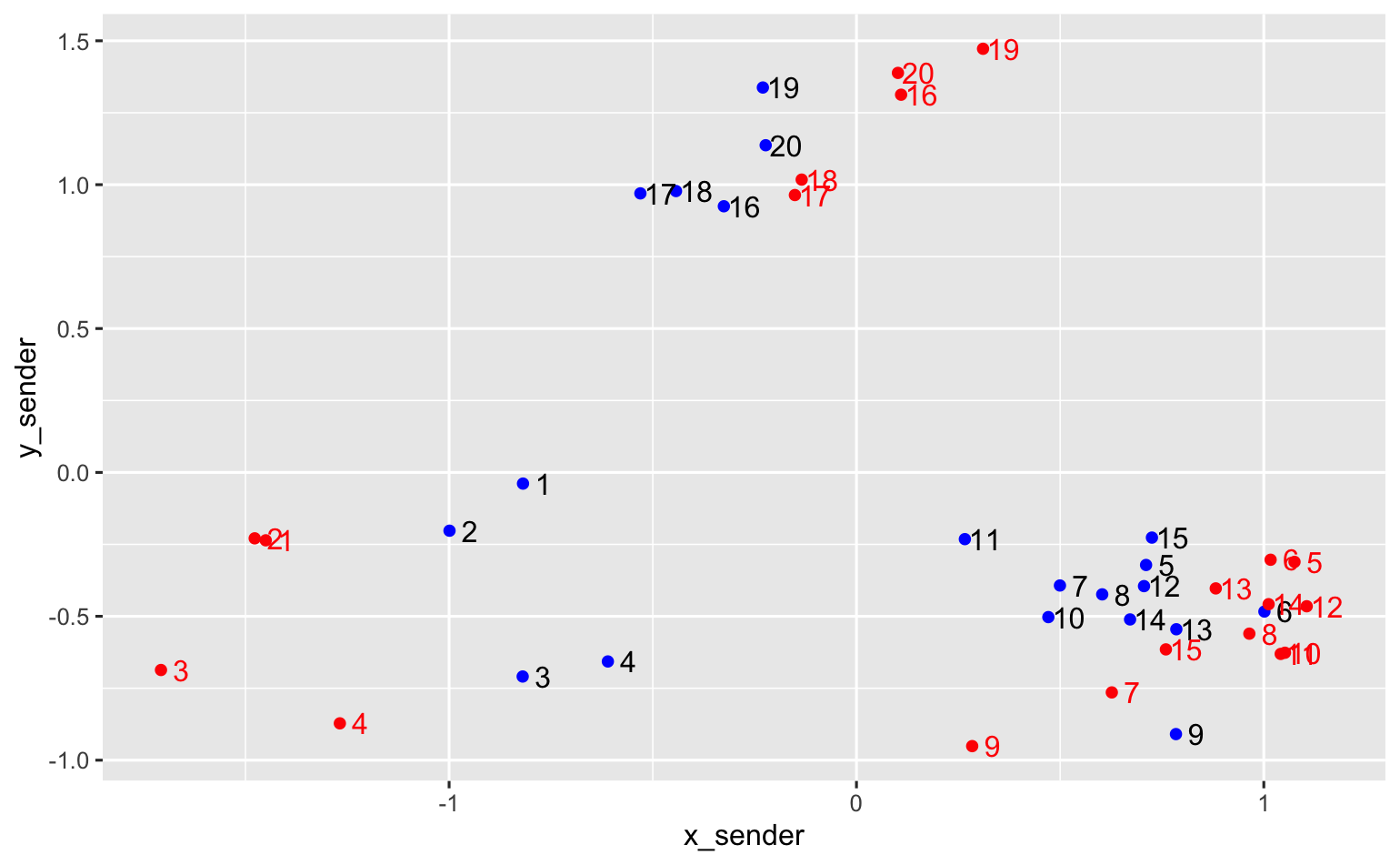

Figure S3: Visualizing the remaining variation left in the factor space for the simulated data with an unmeasured variable. Blue dots are senders, and red dots are receivers. Similar distances and directions from the origin (0,0) indicates that those individuals are more likely to interact.
